# Common symbiotic signalling pathway not essential for formation of functional mutualisms with endophytic fungi

**DOI:** 10.1101/2025.01.10.632311

**Authors:** Alex Williams, Besiana Sinanaj, Victor Rodriguez-Morelos, James Prout, Nathan Howard, Emily Durant, Silvia Pressel, Katie J. Field

**Affiliations:** Plants, Photosynthesis and Soil, School of Bioscience, University of Sheffield, Sheffield, S10 2TN, UK; Department of Biochemistry and Metabolism, John Innes Centre, Norwich Research Park, Norwich, NR4 7UH, UK; Department of Life Sciences, Natural History Museum, London, SW7 5BD, UK

**Keywords:** symbiosis, plant-soil interactions, mycorrhiza, Mucoromycotina ‘fine root endophyte’, common symbiotic signalling pathway

## Abstract

Most plants form mutualistic symbioses with soil fungi, including arbuscular mycorrhizal (AM) fungi. These fungi usually transfer soil nutrients to plants and assimilate carbon from host plant photosynthesis. Recently, Mucoromycotina ‘fine root endophytes’ (MFRE) were identified as nutritionally mutualistic and widespread fungal symbionts of plants, establishing MFRE as a new class of mycorrhizal fungi. However, the regulatory mechanisms for MFRE symbioses are completely unknown. Other symbionts, like AM fungi, use the ‘Common Symbiotic Signalling Pathway’ (CSSP) to establish symbiosis. To explore whether MFRE interactions also involve this pathway, we cultured MFRE with CSSP mutants of *Medicago truncatula* which show impaired AM symbioses and tracked carbon and nutrient transfers using isotope tracers. Results show no differences in root colonization or nutrient exchange, suggesting MFRE symbioses are regulated by different molecular mechanisms. This finding highlights the unique nature of MFRE symbiosis, broadening our understanding of diverse fungal symbioses and their evolutionary significance.

## Introduction

Mycorrhizal associations in plants, and mycorrhizal-like associations in non-vascular plants, are near-ubiquitous mutualistic symbiosis between certain groups of soil fungi and most land plant phyla (Brundrett and Tedersoo, 2018). Through specialised structures, including arbuscules and extraradical mycelium, mycorrhizal fungi assimilate and transfer soil nutrients such as phosphorus (P) and nitrogen (N) to their host plant, while the plant passes photosynthesis-derived carbon resources to the fungus in the form of sugars and lipids (Roth and Paszkowski, 2017). This bi-directional exchange of plant-fixed carbon for fungal-acquired nutrients is the essential characteristic of mycorrhizal symbioses (Strack et al., 2003; Brundrett, 2004; Smith and Read, 2010)

Plant-fungal symbioses have evolved over at least 460 million years (Strullu-Derrien et al., 2018). It has long been hypothesised that by enhancing plant access to essential mineral nutrients, mycorrhizal-like fungi played a pivotal role in facilitating the colonisation of the terrestrial environment by plants during the Cambrian period (Pirozynski and Malloch, 1975). This hypothesis is particularly pertinent when considering the absence of roots in early land plants (Kenrick and Strullu-Derrien, 2014) together with the plant-inaccessible, nutrient-poor nature of the skeletal, mineral soils of that time (Xue et al., 2023) as well as evidence of lipid transfer to AM partners in the primitive land plant Marchantia paleacea, that is conserved in higher land plants but absent in algae (Rich et al., 2021). Further evidence supporting the crucial role of symbiotic fungi in plant terrestrialisation has been accumulating over the last 50 years across various disciplines and sources, including fossil (Remy et al., 1994; Taylor et al., 1995; Dotzler et al., 2006), physiological (Humphreys et al., 2010; Field et al., 2012; Field et al., 2015) and molecular analyses (Simon et al., 1993; Redecker et al., 2000; Wang et al., 2010; Oldroyd, 2013; Delaux et al., 2013). The ability of plants to associate with certain microbial symbionts has largely been explained by the existence of a suite of highly conserved molecular mechanisms, known as the Common Symbiotic Signalling Pathway (CSSP (Oldroyd, 2013; Genre et al., 2020)). The widespread conservation of CSSP elements indicates an ancient and critical role for mycorrhiza-forming fungi during the land-expansion of plants. The genes that make up the CSSP are not only largely conserved across modern land plant lineages (Radhakrishnan et al., 2020), but some of these genes are also present in charophytic algae, the closest green algal relative to land plants (Delaux et al., 2015). Widespread conservation of the CSSP across the land plant phylogeny indicates it is still beneficial to most plants, where its function is thought to be key for the establishment and regulation of mycorrhizal symbioses.

To date, the dominant focus of research has centred on arbuscular mycorrhizal (AM) fungi, neglecting the full scope of endophytic, root-fungal interactions that also commonly occur, including those formed by Mucoromycotina ‘fine root endophytes’ (MFRE (Orchard et al., 2017; Hoysted et al., 2019)). MFRE are a group of soil fungi belonging to the sub-phylum Mucoromycotina, parallel to AM-encompassing Glomeromycotina fungi (Bidartondo et al., 2011; Spatafora et al., 2016; Chang et al., 2022; Zhao et al., 2023), in the phylum Mucoromycota. Recently, MFRE have emerged as being physiologically, ecologically, and evolutionarily important root fungal endophytes across a wide variety of plants and ecosystems (Hoysted et al., 2018), forming mycorrhizal associations with species from most land plants (Hoysted et al., 2019; Hoysted et al., 2021; Hoysted et al., 2023). Fossil evidence and molecular dating of the divergence of Glomeromycotina from Mucoromycotina suggest plant associations with MFRE are likely to be at least as ancient as AM fungi associations, with some Rhynie Chert fossils providing glimpses of MFRE-like fungal structures within early land plant tissues (Southworth, 2012; Strullu-Derrien et al., 2014). The full range, frequency and ecological relevance of plant associations with MFRE in modern flora remain to be fully explored, but it has become increasingly apparent that these associations are widespread, abundant, and have persisted across evolutionary timescales (Hoysted et al., 2018).

While new understanding about the functional, ecological, and evolutionary significance of MFRE associations is emerging, key outstanding questions regarding the mechanisms controlling their formation and function remain (Prout et al., 2024). The genes underpinning the CSSP facilitate and regulate processes for mutual recognition, hyphal colonisation and tissue ingress, arbuscule formation, and carbon for nutrient exchanges in mycorrhizal symbioses (Oldroyd, 2013). Symbiosis begins with mutual recognition of host-plant and microbe, via root exuded strigolactones (Lanfranco et al., 2018) and microbial signals (myc or NOD factors, *e.g.* lipo-chitooligosaccharides (Zipfel and Oldroyd, 2017)). A suite of specific genes and processes occur (Bravo et al., 2016; Radhakrishnan et al., 2020), that are broadly homologous among host plants, to mediate interactions with AM fungi but also other symbiotic rhizobacteria, such as nitrogen-fixing *Rhizobia* G. and *Pseudomonas fluorescens* (Sanchez et al., 2005). Extensive characterisation of various aspects of the CSSP have been performed in the model legume *Medicago truncatula* (Oldroyd, 2013), establishing a sequential interaction and signalling cascade that leads to the successful formation of root symbioses with beneficial microbes. In *M. truncatula*, initial mycorrhization is defined by a perinuclear spike in Ca^2+^ activated via a signalling cascade under the control of the receptor kinase Mt*DMI2*. Subsequent activation of the Ca^+2^/calmodulin dependent kinase Mt*DMI3* leads to a decoding of Ca2+ spiking, and regulating symbiotic transcription factors essential for either nodulation or arbuscule formation (Tian et al., 2020). In AM symbiosis the activation of Mt*DMI3* indirectly regulates a series of transcription factors, including the master regulator *RAM1* (a member of the *GRAS* transcription factor family), required for development of a nutritionally functional mutualism. Although not a process shared with rhizobial nodulation, *RAM1*-dependent intracellular arbuscule development (Gobbato et al., 2012) acts as the primary site of C-for-nutrient exchange, with the aid of various downstream transcription factors including *WRI5*, essential for lipid transfer (Jiang et al., 2018).

Respectively, CSSP gene mutants (*dmi2*, *dmi3* and *ram1*) in *M. truncatula* are all impaired in functional AM symbiosis (Wais et al., 2000; Rich et al., 2015). However, in MFRE the development of arbuscule-like structures may not be necessary for the formation of a functional relationship (Hoysted et al., 2019; Hoysted et al., 2021; Hoysted et al., 2023) and their presence is rare in axenically cultured symbioses (Prout et al., 2024). This suggests the bi-directional transfer of resources between plants and MFRE may occur through inter-or intraradical hyphal mycelium and/or at hyphal tips. It might also indicate transfer of a wider range of C-based molecules from host to fungus than are typical in AM symbioses, particularly given the facultative biotrophic and potential saprotrophic capabilities of MFRE (Field et al., 2015; Hoysted et al., 2023). As such, considering the phylogenetic distance between MFRE and AM fungi as well as the relatedness of MFRE to saprotrophs and pathogens (Chang et al., 2022), we hypothesise that late-stage mechanisms of the CSSP which regulate arbuscule formation may not be operational within plant-MFRE interactions. Instead, MFRE may colonise hosts through an entirely different pathway to those described in the CSSP. If this is the case, we expect *M. truncatula* CSSP mutants impaired in their ability to form and maintain AM symbioses to retain the ability to maintain formation of functional symbiosis with MFRE. Should the CSSP prove critical for establishment, regulation, and function of plant-MFRE symbiosis it would provide further evidence that the CSSP operates as a fundamental mechanism for symbiosis across mycorrhizal lineages (Prout et al., 2024). Should the CSSP prove unnecessary for the formation and function of MFRE-plant symbioses, it would suggest these interactions operate outside of the bounds of the general framework for mycorrhiza symbioses, indicating a separate evolutionary history as ancient as Glomeromycotina AM symbioses. The presence of an alternative mechanism of mycorrhizal symbiosis, persistent across the time-scale over which AM’s have evolved (Zhao et al., 2023), would challenge our understanding of plant evolutionary history and terrestrialisation.

To address this fundamental knowledge gap, we used radioactive and stable isotope tracers and metabolomics analysis to investigate the capacity of MFRE to form nutritionally mutualistic associations with *M. truncatula* mutants deficient in critical elements of the CSSP.

For methodology see **Methods File 1**

## Results

### Plant biomass and root colonisation by MFRE are unaffected by mutations in the CSSP

Root colonisation and extraradical mycelial development was extensive in all MFRE-inoculated plants, with intracellular structures and morphology consistent with that described previously in other MFRE-angiosperm symbioses (Fig. 1a-b (Hoysted et al., 2023)). There were no differences in the types of structures observed in WT or CSSP mutant plant roots, including fine hyphae, fan-like branching hyphae, ‘;vesicles’ or swellings, and hyphal coils typical of vascular plant-MFRE colonisations (Fig. 1 b (Hoysted et al., 2019; Hoysted et al., 2023)), but the abundance of these structures was lower in *dmi2-1* and *dmi3-1* (Fig. 1 d; ANOVA; F_3,77_=6.16, *P*<0.001) and the extent of hyphal colonisation was lower in *dmi3-1* (Fig. 1 c; ANOVA; F_3,77_=3.35, *P*=0.023) compared to WT (See Fig. S1 for more images). Differences in colonisation did not relate to growth as there were no differences in shoot biomass (Fig. S2) between the wild-type A17, CSSP mutants or the non-fungal controls (ANOVA, F_3,36_=2.17, *P*=0.11). However, root biomass was larger in *dmi3-1* compared to *dmi2-1* but not the other groups, and none differed from non-fungal controls (ANOVA, F_4,35_=2.78, *P*=0.042). Total biomass was broadly similar across genotypes, suggesting no overall growth promotion of *M. truncatula* by MFRE.

**Figure 1.**
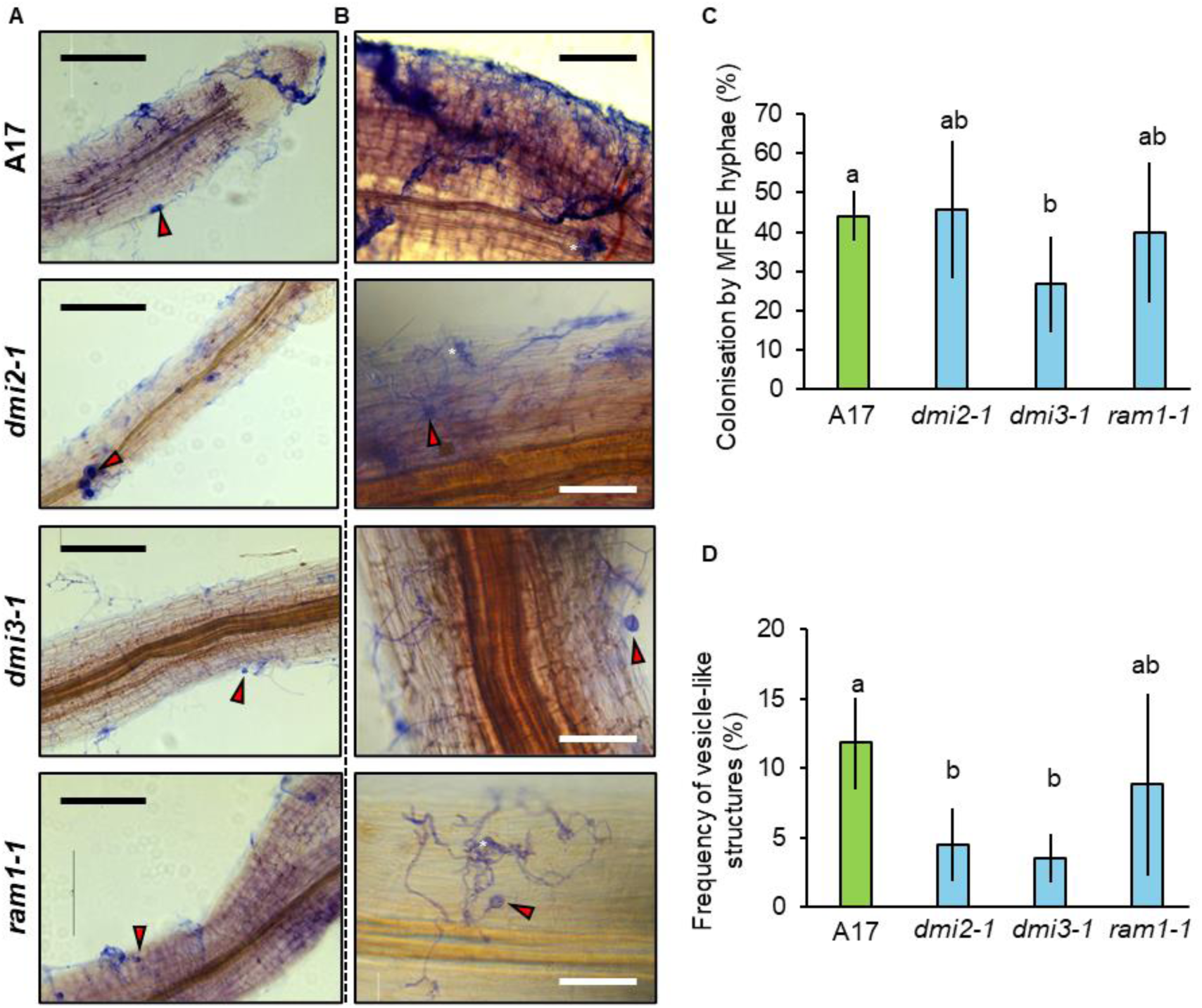
Colonisation of *Medicago trucatula* roots by Mucoromycotina fine root endophytes (MFRE). **(a)** Axenic culture of ink-stained roots of *M. trucatula* and MFRE after 12 weeks co-culture, including wild-type A17 and the mutants *dmi2-1*, *dmi3-1* and *ram1-1*. Fine, irregularly branching MFRE hyphae with large vesicles are shown (red arrows). Mycelium covering the root surface are enlarged in **(b)**, see also Fig. S1); note hyphal clusters (*). Extent of colonisation of *M. trucatula* roots by hyphae **(c)** and characteristic hyphal swellings **(d)**. Letters indicate statistical separation (ANOVA and Tukey’s HSD). Scale bars: **(a)** 250 μm **(b)** 100 μm.

### MFRE colonisation enhances host plant nutrient uptake and is not impaired by CSSP mutations

Total P (plant + MFRE-acquired) concentration (Fig. 2 a) in plant shoots differed between genotypes and non-fungal controls, with shoot [P] of in MFRE colonised plants of all genotypes being greater than non-fungal plants. However, due to large variability in non-inoculated controls, this pattern was borderline non-significant (ANOVA, F_4,35_=2.47, *P*=0.06). MFRE-mediated shoot ^33^P concentration was higher in MFRE-colonised plants than in non-fungal controls, although the differences were not statistically significant (Fig. 2 b; ANOVA, F_4,35_=2.15, *P*=0.09). The *ram1-1* mutant contained the lowest overall ^33^P shoot concentration, consistent with *ram1-1* also having the lower total P values compared to the other genotypes. Neither total P per plant nor total ^33^P accumulation per plant were significantly different between treatments, regardless of host plant genotype (Fig. S3 a-b; ANOVA, F_4,35_=0.93, *P*=0.46; ANOVA, F_4,35_=2.03, *P*=0.11).

**Figure 2.**
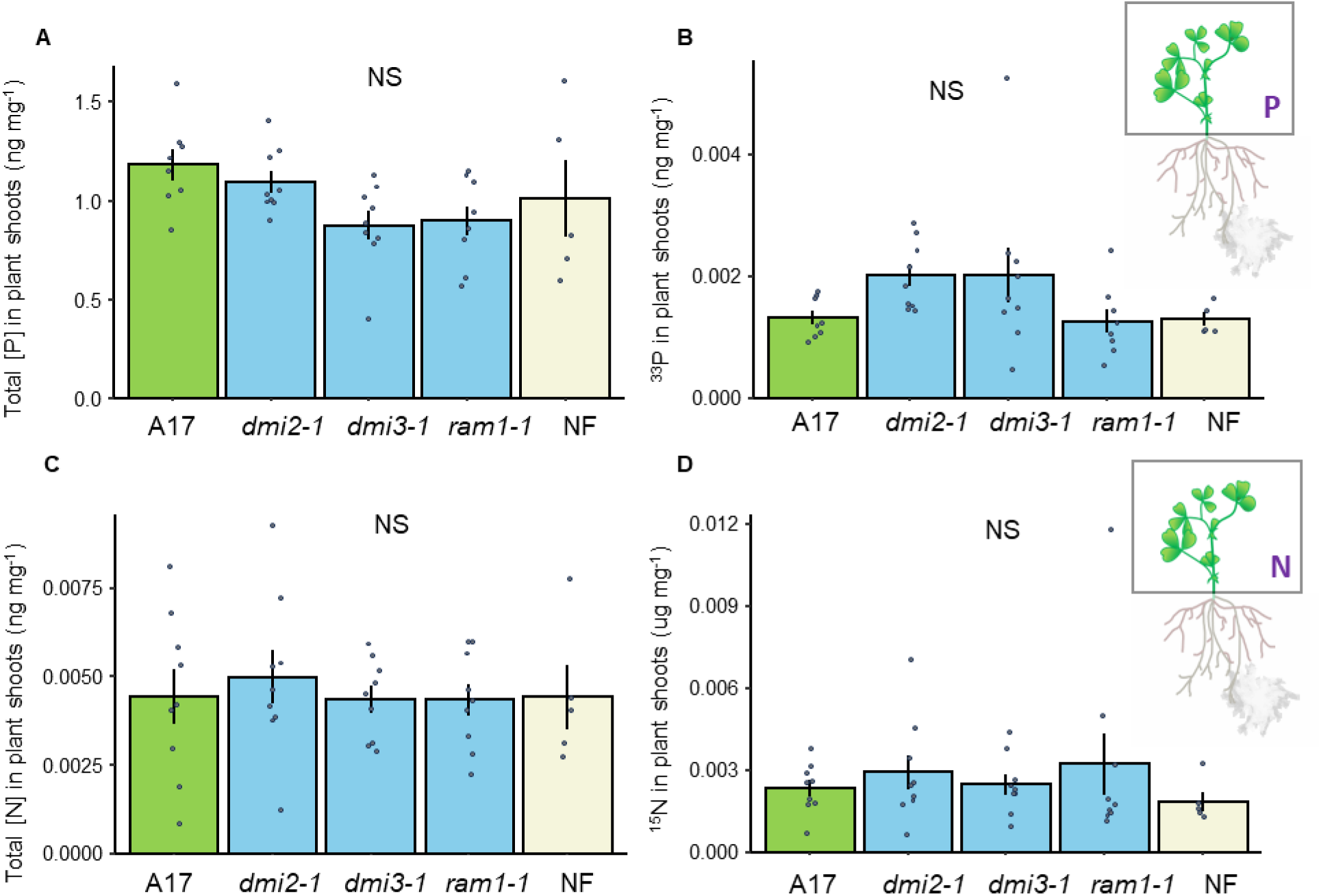
Phosphorus and Nitrogen exchange between *Medicago truncatula* and Mucoromycotina fine root endophytes (MFRE). **(a)** Shoot concentration of phosphorus (ng mg^-1^) in sterile monoxenic cultures with intact fungi across wild-type A17 and the mutants *dmi2-1*, *dmi3-1* and *ram1-1* and non-fungal (NF) control; and **(b)** shoot concentration of ^33^P (ng g^-1^). **(c)** Shoot concentration of nitrogen (ng g^-1^) and **(d)** shoot concentration of ^15^N (ng g^-1^) Represented by box and whiskers are the median (black band), quartiles (ends of rectangle) and extremes (end of bars). Outliers are highlighted with red points. n = 8 for monoxenic cultures with MFRE and n = 5 for non-fungal control.

Total N concentrations in shoots were generally higher in MFRE-colonised plants compared to non-fungal plant shoots, although this was not significant (Fig. 2 c; ANOVA, F_4,35_=0.47, *P*=0.76). Similarly, MFRE-acquired [^15^N] shoot concentrations were higher for plants colonised by MFRE than non-fungal plants, albeit the difference was not statistically significant (Fig 2 d; ANOVA, F_4,36_=0.50, *P*=0.73). Concentrations of MFRE-acquired ^15^N were the lowest in *ram1-1* and highest in *dmi2-1* shoot tissues comparted to the WT (Fig. 2 d). Neither total N per plant nor total ^15^N accumulation per plant were significantly different between treatments, regardless of host plant genotype (Fig. S3 c-d; ANOVA, F_4,35_=0.36, *P*=0.83; ANOVA, F_4,36_=0.32, *P*=0.81).

### Host plants allocate recent photosynthates to MFRE regardless of CSSP mutations

WT, *dmi2-1*, and *ram1-1* plants all allocated recently fixed C to MFRE partners with more plant fixed C being present in the MSR medium containing MFRE extraradical mycelium than in non-fungal plants (Fig. 3 a; ANOVA, F_4,34_=4.00, *P*<0.01). There was no significant difference in the amount of plant-fixed C transferred to MFRE between WT (A17), *dmi2-1* and *ram1-1* genotypes, while C transfer from plant to MFRE was not observed in *dmi3-1* plants. In terms of percentage of C fixed by photosynthesis during the labelling period which was exported to the MFRE/ MSR medium, we observed a similar pattern, although, in contrast to the values for total C, the differences were not significant (Fig. 3 b; ANOVA, F_4,34_=1.51, *P*=0.22).

**Figure 3.**
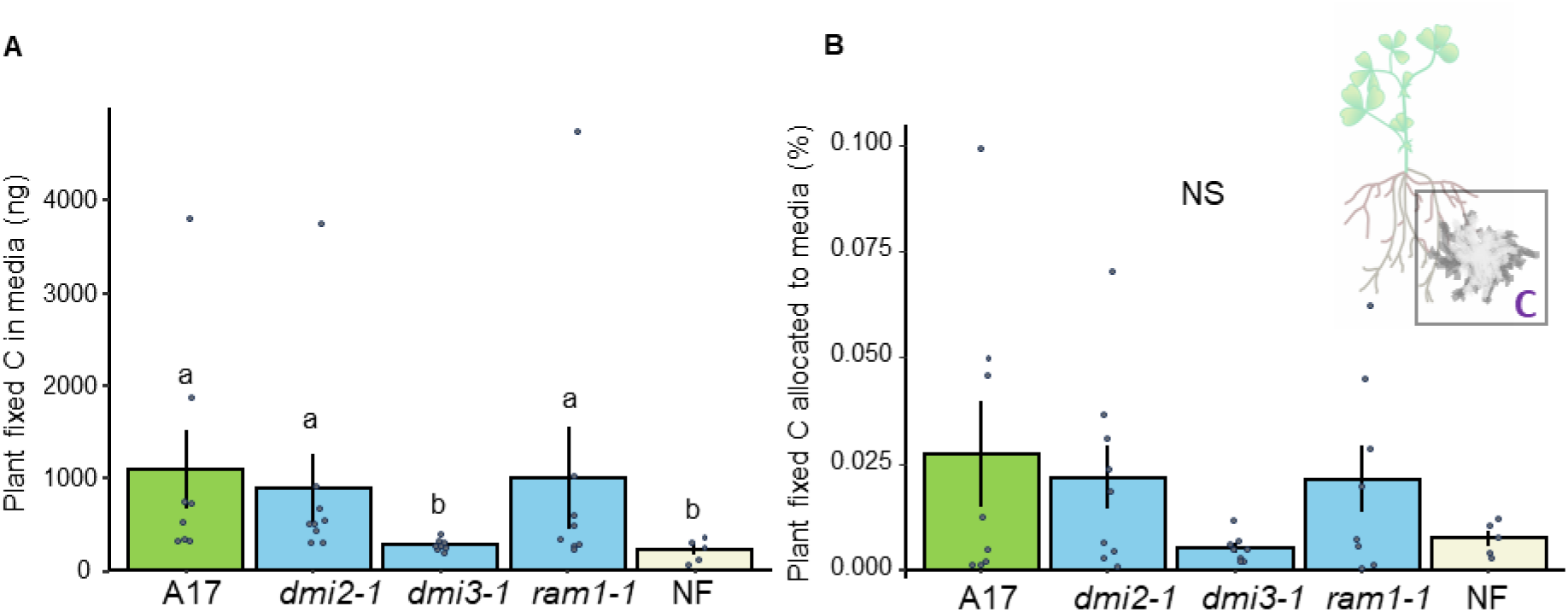
Total plant-derived carbon present in Mucoromycotina fine root endophytes (MFRE) extraradical mycelium. **(a-b)** Total plant-derived carbon present in media and MFRE extraradical mycelium across *Medicago truncatula* wild-type A17 and the mutants *dmi2-1*, *dmi3-1* and *ram1-1* with MFRE fungi present or plant-only microcosms where no MFRE were present (NF) after a 24-h labelling period with radio-isotope ^14^C. **(a)** Mean total plant fixed C in MFRE hyphae within media (ng) and **(b)** mean percentage allocation of ^14^C to the MFRE medium are displayed. Letters indicate statistical differences where *p*<0.05 (ANOVA and post-hoc analysis). Bars represent standard error. n = 8 for microcosms with MFRE present and n = 5 for NF.

### Evidence of fatty acid biosynthesis in extraradical MFRE mycelium

Untargeted analysis of metabolomic profiles showed strong influence of tissue type and MFRE colonisation status in both ionisation modes on plant metabolism (Fig. 4 a & b) suggesting both local and systemic impacts of colonisation by MFRE. Focusing on local signals of colonised and non-fungal root material, there were a small number of significantly altered metabolites (38 enriched metabolites in ESI-and 55 in ESI+ with 24 and 22 enriched in NF respectively; Fig. S4 a-b). These metabolites were predominantly lipid and terpenoid metabolites, but there were also many that were unidentified (29.7 %; Fig. S4 e; Table S2). To gain insight into the systemic signature of MFRE colonisation we also looked at host plant shoot material, identifying 40 metabolites enriched in ESI-and 69 enriched in ESI+, with 56 and 52 enriched in non-fungal plant shoot material respectively (Fig. S4 c-d). Many of those enriched in MFRE colonised plants were lipids (Fig. S4 e), suggesting colonisation by MFRE induced a systemic production of C-rich compounds.

**Figure 4.**
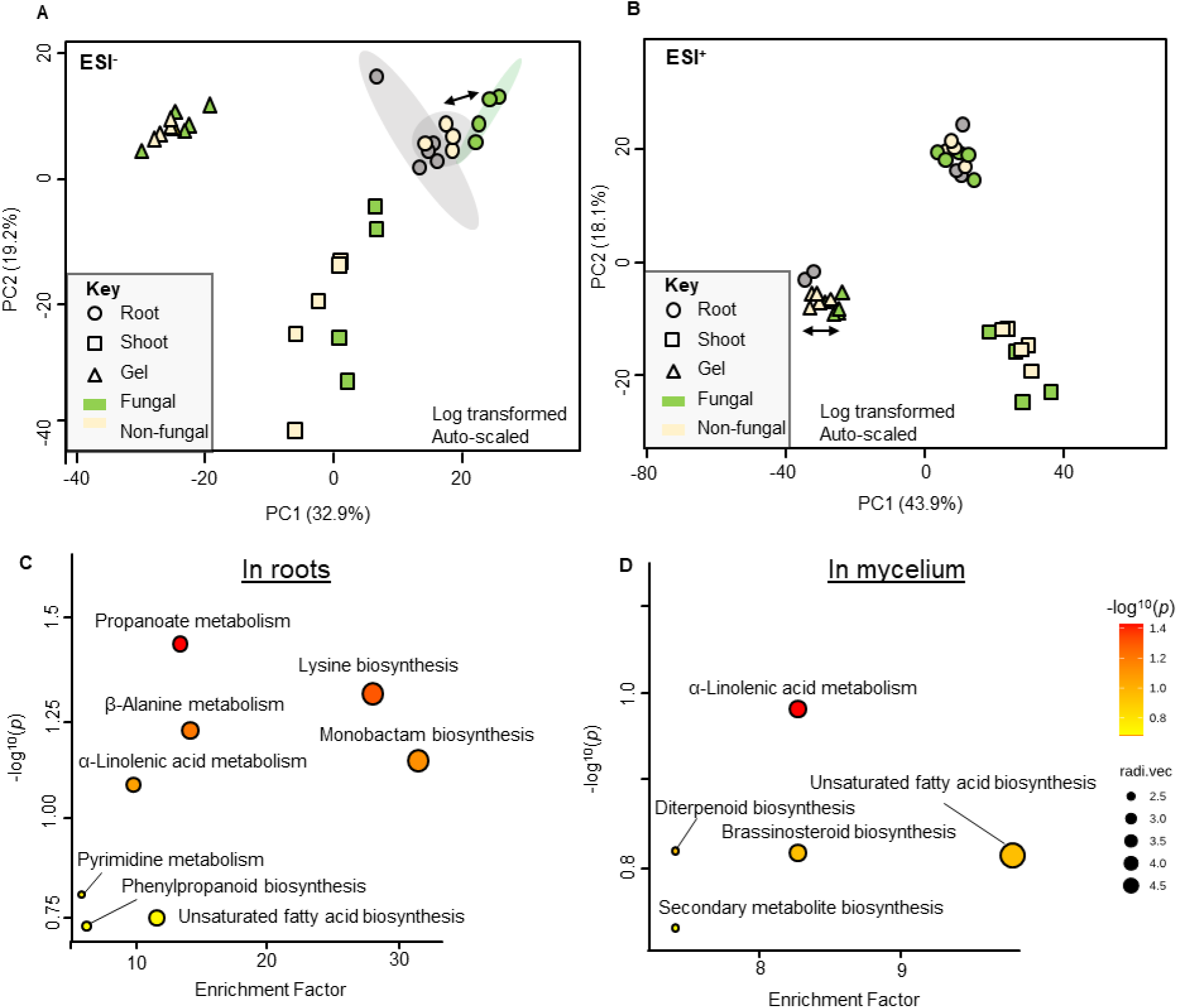
Principal Component Analyses of metabolomic profiles of MFRE colonised *Medicago truncatula* in ESI-**(a)** and ESI+ **(b)** ionisation modes, among sample types tested (shoot, root and media) of colonised (green) and non-colonised (cream) plants. Roots distal to those colonised were also tested (grey circles). Pathway analysis (mummichog) illustrate the top differentially affected pathways in roots **(c)** and MFRE extraradical mycelium **(d)** of ESI-profiles. Richness of red indicates significance and size indicates enrichment factor.

For a more parsimonious indication of pathways that were affected by MFRE colonisation, we employed Mummichog analysis on the ESI-root samples. We identified a number of pathways highly altered in colonised roots, including lysine and β-alanine biosynthesis along with phenylpropanoid metabolism, fatty acid biosynthesis and propanoate metabolism (among other pathways; Fig. 4 c, Table S3). As these annotations cannot distinguish between plant-and fungal-derived metabolites, we also analysed MSR medium containing only MFRE extraradical mycelium to identify pathways that were likely fungal-derived. Here we identified secondary metabolite biosynthesis, brassinosteriod and diterpenoid biosynthesis and fatty acid biosynthesis, including α-linolenic acid metabolism (Fig. 4 d).

## Discussion

We tested whether the CSSP, a molecular pathway shared with other microbial nutritional mutualists such as AM fungi and *Rhizobia* (Oldroyd, 2013), was important in the establishment and function of MFRE symbiosis in the model legume, *M. truncatula,* using mutants defective in critical CSSP elements. Our findings reveal that two of the mutants tested (*dmi2-1, ram1-1*) were largely unaffected in the extent of root colonisation by MFRE hyphae, the types of fungal structures present within colonized roots, and the bidirectional exchange of plant-fixed C for MFRE-acquired nutrients compared to colonised WT (A17) plants. However, mutant *dmi3-1* plants tended to have a lower percentage of root colonisation and greatly reduced C allocation to the fungus than WT or the other mutants tested.

CSSP genes *DMI2* and *RAM1* are centrally important to early and late stages of AM symbiosis. When *DMI2* is impaired, no intraradical colonisation of host plant roots by AM fungi occurs (Seddas et al., 2009) while RAM1 is required for full development of arbuscules (Pimprikar et al., 2016) which are associated with transfer of plant lipids/fatty acids to AM fungal symbionts. However, in our experiments, MFRE colonisation of *Medicago* roots was not impaired in either *dmi2* or *ram1* mutant plants compared to WT plants. Instead, intra-and intercellular fungal structures including fine branching hyphae with characteristic fan-like branching patterns and vesicle-like structures were present, consistent with typical MFRE colonisation in WT plants. Critically, using radio-and stable isotope tracing, we found no effect of mutations in *DMI2* or *RAM1* on C for nutrient (N and P) exchange – a key diagnostic feature of functional mycorrhizal symbioses - between MFRE and their host plants compared to WT plants. Together, these lines of evidence indicate that the loss of function of *RAM1* or *DMI2* has an inconclusive impact on the establishment, development or functionality of plant-MFRE symbioses. While these genes have an essential role in AM colonisation (Oldroyd, 2013), to our knowledge bi-directional carbon for nutrient exchange between AM fungi and host plants has yet to be demonstrated in CSSP mutants. As we show carbon transfer to MFRE symbionts was maintained comparative to WT plants in the *ram1* and *dmi2* mutants, along with the presence of metabolites associated with fatty acid biosynthesis in the mycelial tissue, our findings could be interpreted that plant-derived C is transferred to MFRE under an alternative dominant form to that which is supplied to AM fungi (Luginbuehl et al., 2017).

*RAM1* regulates *RAM2* and lipid export in *M. truncatula* (Gobbato et al., 2013). Transfer and assimilation of host-derived C from *ram1-1* plants to MFRE was not impaired in our experiments, suggesting that the synthesis, storage and transport of more complex C molecules such as fatty acids and lipids occurs within the fungus itself rather than these compounds being transferred directly or indirectly (e.g. through exudation at the rhizoplane) from host plant to fungal symbiont. Our metabolomics analyses show increased abundance of lipid metabolites and their likely metabolic products (terpenoids), as well as the upregulation of lipid biosynthesis pathways, not only in colonised root material where phytogenesis cannot be ruled out, but critically also in the MFRE extraradical mycelium. AM fungi lack the ability to produce their own lipids as they do not have the required genes to produce the necessary fatty-acid synthases (Tang et al., 2016). As many Mucoromycotina fungi are oleaginous (lipid-accumulating (Kosa et al., 2018), it is possible that plant symbiotic Mucoromycotina, including the Lyc1 MFRE isolate used in our experiments, also possess the ability to synthesise their own complex C molecules through metabolism of more simple carbon molecules, such as hexoses, and subsequent biosynthesis of fatty acids and lipids. The ability to synthesise fatty acids and lipids provides a physiological mechanism that supports the observed facultatively symbiotic lifestyle of plant-symbiotic MFRE (Hoysted et al., 2018; Hoysted et al., 2023; Prout et al., 2024), providing these fungi with a niche and plasticity that is unavailable to obligately biotrophic fungal symbionts which rely entirely on their host plant for their C nutrition.

Despite our findings that *dmi2-1* and *ram1-1* mutants were not impaired in formation or function of nutritional mutualisms with MFRE, *dmi3-1* roots were less colonised by MFRE hyphae than WT plants, with formation of fewer characteristic structures and reduced transfer of plant C to MFRE mycelium. These results strongly support there being a role for *DMI3* in MFRE symbiosis formation, possibly outside of its role in the CSSP. In the CSSP, *DMI3* acts as master regulator for mycorrhization by AM fungi to take place. It is necessary for decoding calcium and calmodulin oscillations which are generated via the sustained activity of *DMI1* and calcium/calmodulin nucleotide-gated channels during the earliest stages of symbiosis (Khatri et al., 2022). Removal of *DMI3* typically results in impaired AM symbiosis and drastically reduced colonisation (Seddas et al., 2009). However, outside the CSSP, *DMI3* and orthologues (CC) have been implicated in the management of several plant-microbe interactions. For instance, DMI3 shows a central role in managing the infection of various pathogens and endosymbionts, where loss-of-function results in compromised immunity against the biotroph *Colletotrichum graminicola*, necrotroph *Phoma medicaginis* and the endosymbiont *Oidiodendrum maius* (Genre et al., 2020). In the case of interactions with the beneficial rhizobacteria *Pseudomonas simiae* (previously *fluorescens*) *DMI3* alone was activated during infection while other CSSP genes were not (Sanchez et al., 2005). Together, these studies suggest a regulatory role for *DMI3* in microbial immune management outside of CSSP. There was a general downregulation of immune metabolite pathways relating to immune regulation (including alkaloids and flavonoids) in our MFRE-colonised plants. Although further evidence, considering different alleles and genetic background would help to verify its role, it is tempting to speculate that during MFRE-plant symbioses *DMI3* may act as a master-regulator for general immune function via some alternative pathway.

Although our findings have far-reaching consequences, our experiential design is somewhat limited in terms of ecological relevance due to exclusion of other interacting microbes that would typically be present within the rhizosphere. However, this absence of biotic interactions permits mechanistic insights to be made without the confounding effects of other organisms. Our data displays variability within treatments due to the nature of isotope tracer studies with living organisms in monoxenic culture which inevitably produce variation (e.g. (Hoysted et al., 2023; Howard et al., 2024), and the relatively short timeframe of the labelling period (24 hr). This is partly due to the limitations of the experimental design, such as the risk of contamination by other interacting microbes, and limited growth space within microcosms for the plants which constrains the duration of incubation with tracers. Furthermore, there is a pressing need to evaluate whether the observed phenotypes are consistent across different MFRE species, as new isolates are found, and across different plant functional groups. Future experiments are also needed to investigate the relevance of the mechanisms investigated here in soil-based systems replete with natural rhizosphere microbial communities over longer time scales.

While the apparent lack of function of the CSSP genes *DMI2* and *RAM1* in establishing functional plant-fungal symbioses with MFRE challenges the assumed similarities between MFRE and AM fungal symbioses, it also reveals fascinating details about this unique partnership. Unlike AM symbioses, where arbuscule formation (regulated by components of the CSSP) is necessary for lipid transfer, MFRE appear to not require such specialised structures. This, together with evidence from our isotope tracing and metabolomics analyses, suggests a simpler exchange mechanism to be functional in MFRE symbioses, potentially involving the transfer and utilisation of simple molecules, such as hexoses, to the fungus whereupon the biosynthesis of more complex C-based molecules occurs. Furthermore, considering the presence of fine root endophytes in fossilized plants, such as Horneophyton (Strullu-Derrien et al., 2014) and the potentially earlier divergence of Mucoromycotina (MFRE fungal lineage) compared to Glomeromycotina (AM fungal lineage (Bidartondo et al., 2011; Wang et al., 2023), the lack of response to RAM1 mutation in host plants by MFRE points towards the existence of alternative evolutionary pathways for plant-fungal symbioses. Although currently inconclusive, these findings question the existence of a single, conserved genetic pathway for establishment, development and function of all endomycorrhizal symbioses. Instead, they nod toward a broader view where alternative mycorrhizal symbiotic options were available and likely important in the emergence and establishment of a mycorrhizal terrestrial land flora (Field et al., 2015). The specific mechanisms of the CSSP may be less crucial for MFRE function than that of AM, opening doors for exploring alternative pathways in, and the importance of, non-AM nutritional mutualisms formed between plants and fungi.

## Acknowledgements

We thank Dr Heather Walker for (University of Sheffield) for mass spectrometry analysis of ^15^N samples. KJF, AW, BS, NH, SP and VRM are supported by the ERC CoG “MYCOREV” (865225). ED is supported by the NERC ECORISC DTP, JP is supported by a University of Sheffield PhD studentship. We thank the De Laszlo Foundation for generously supporting PhD student research at the University of Sheffield. We thank Giles Oldroyd and Tom Thirkell (CropScienceCentre, Cambridge) for providing *M. truncatula* mutant seeds and discussion of our findings.

## Author Contributions

AW and KJF conceived and designed the investigation. AW, NH, JP and ED undertook the experiments, SP and VRM provided fungal isolates. AW, NH, ED, JP and BS conducted the tracing experiment and collected data. AW analysed the results. AW, JP, KJF and SP interpreted and discussed the results. AW led the writing; all authors discussed results and commented on the manuscript. AW and KJF agree to serve as authors responsible for contact and ensure communication.

## Supplementary Materials

**Figure S1.**
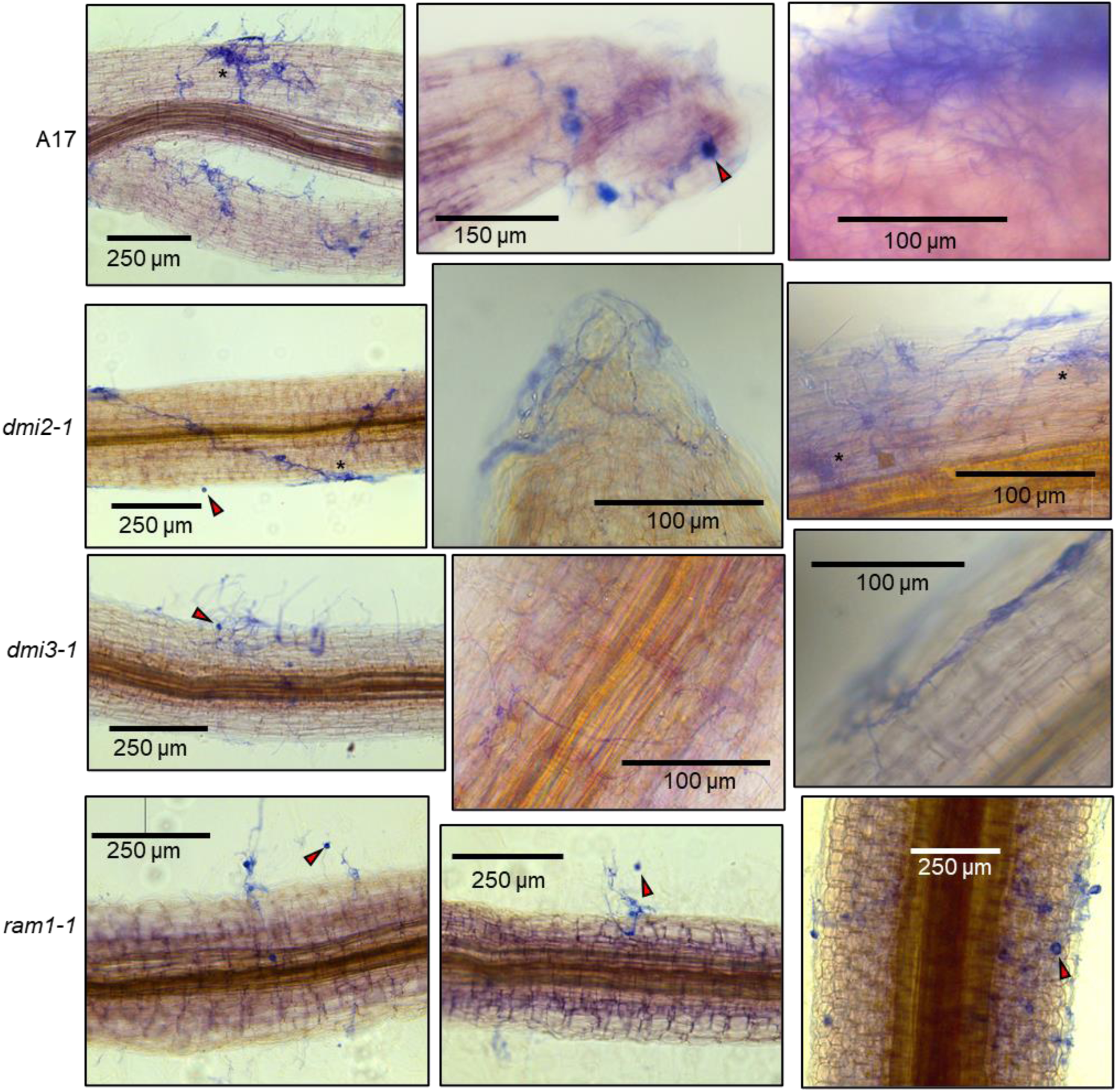
Colonisation of *Medicago trucatula* roots by Mucoromycotina fine root endophytes (MFRE). Axenic culture of ink-stained roots of *M. trucatula* and MFRE after 12 weeks co-culturing. Four genotypes are displayed wild-type A17 and the mutants *dmi2-1*, *dmi3-1* and *ram1-1*. Fine, irregularly branching hyphae with large vesicles (red arrows) and hyphal clusters (asterix) are shown. Note abundant mycelium covering the root surface. Scale bars are all indicated on the images.

**Figure S2.**
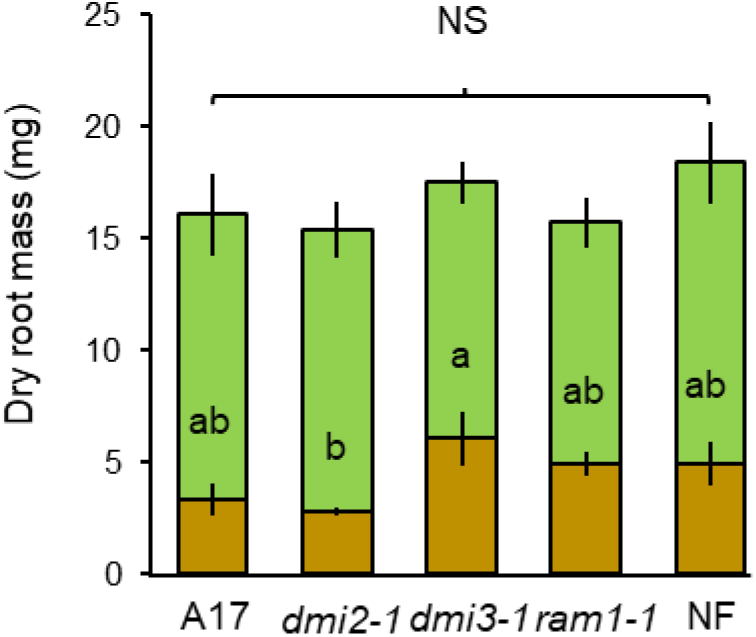
Biomass measurements of *Medicago trucatula* genotypes wild-type A17, *dmi2-1*, *dmi3-1* and *ram1-1*, or a non-fungal control (NF). Bottom bars (brown) represent root biomass (mg), top bars (green) represent shoot biomass. Letters indicate statistical differences in root biomass (ANOVA and post-hoc analysis).

**Figure S3.**
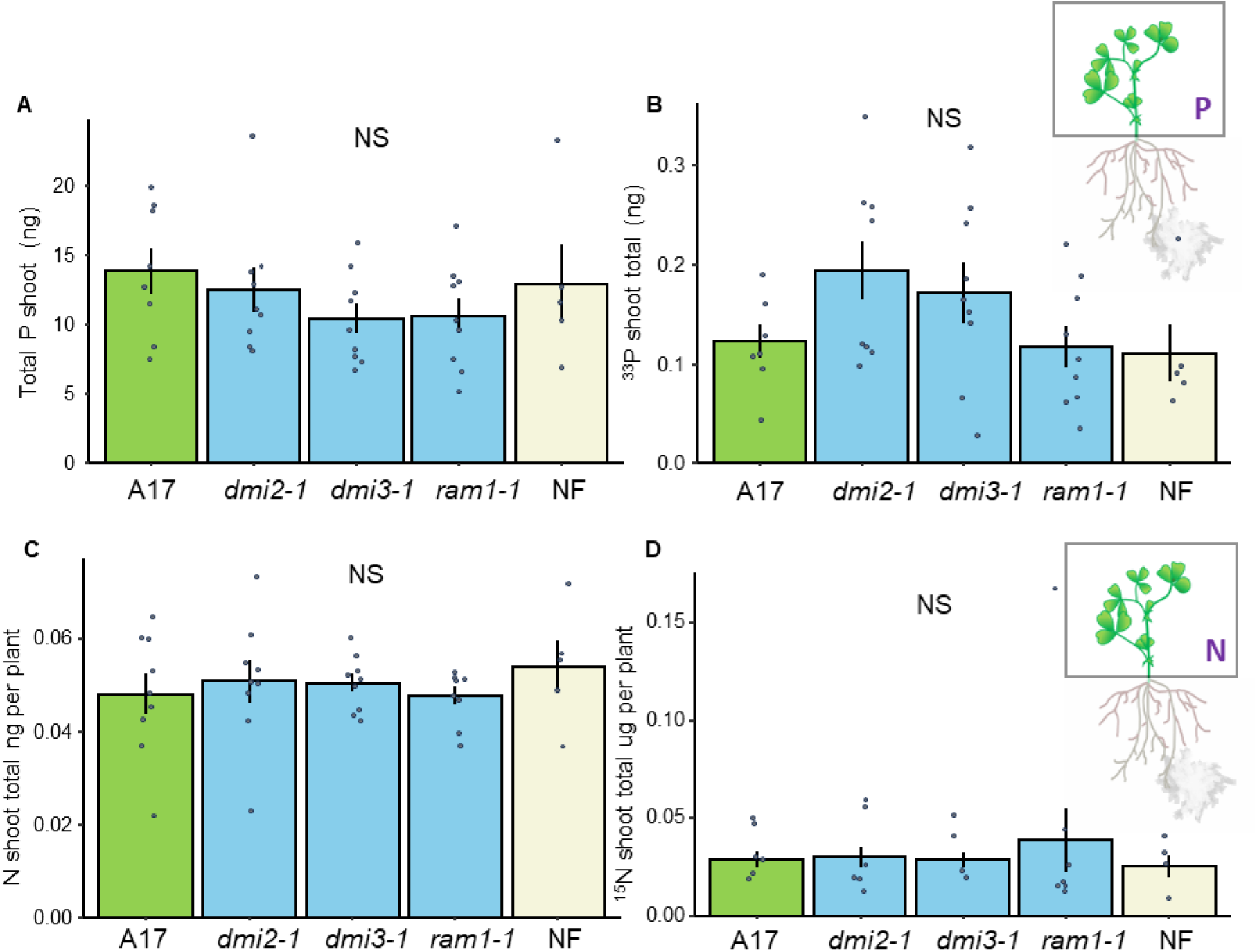
Total Phosphorus and Nitrogen exchange (per plant) between *Medicago truncatula* and Mucoromycotina fine root endophytes (MFRE). **(a)** total shoot tissue phosphorus (ng) in sterile monoxenic cultures with intact fungi across wild-type A17 and the mutants *dmi2-1*, *dmi3-1* and *ram1-1* plus a non-fungal control and **(b)** total shoot fungal-acquired ^33^P in *M. trucatula* **(c)** total shoot tissue nitrogen in the same genotypes (ng). **(d)** total fungal-acquired ^15^N in M. trucatula shoot. n = 8 for monoxenic cultures with fungi and n = 5 for no fungal control.

**Figure S4.**
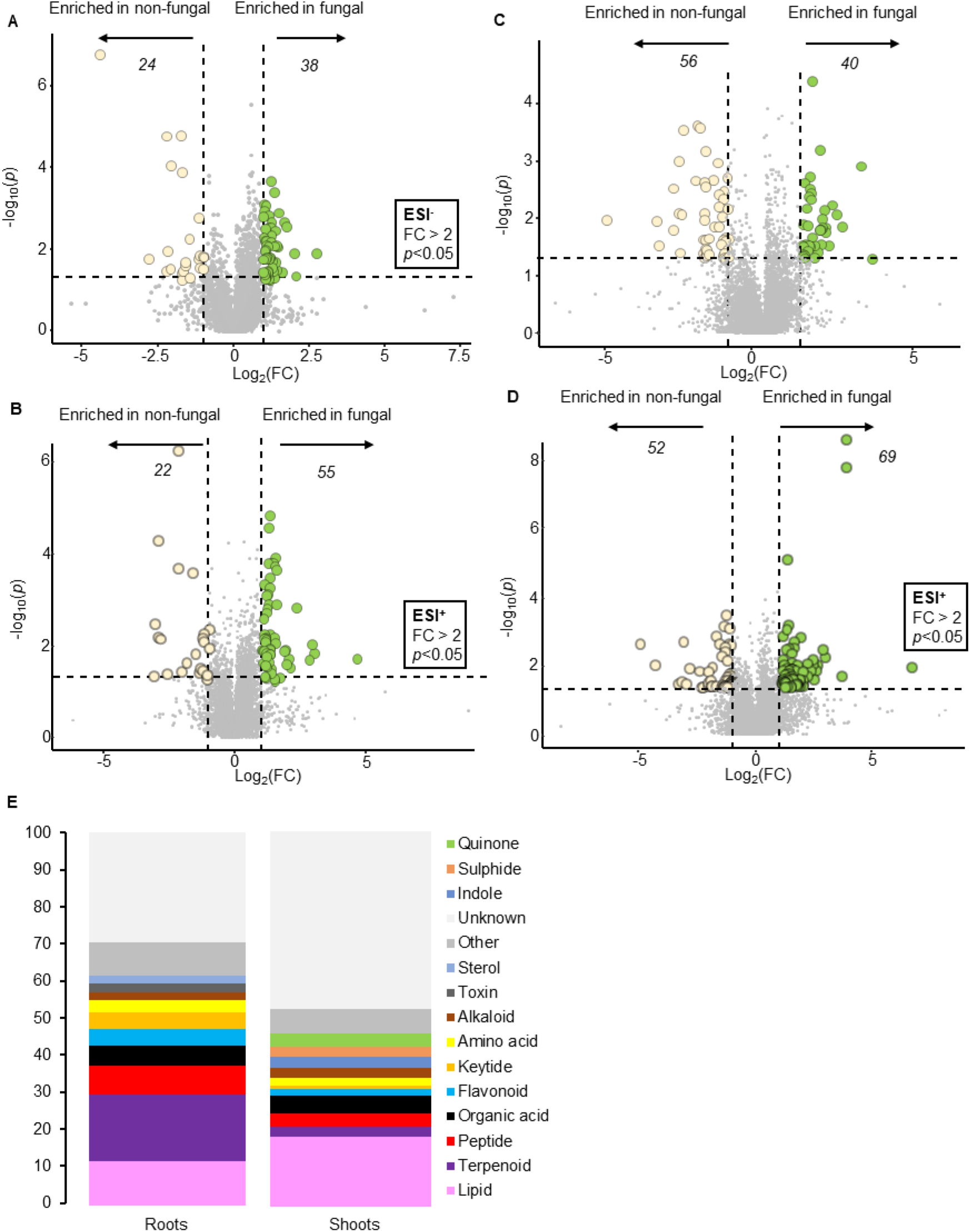
Volcano plots of root material either with or without MFRE. Shown are metabolites enriched at P<0.05 with a FC of 2 in fungal (right, green circles) or non-fungal (left, cream circles). Shown is root material in **(a)** ESI^-^ and **(b)** ESI^+^ mode. Shoot material for **(c)** ESI^-^ and **(d)** ESI^+^ are also shown. Data were log-transformed and auto-scaled, consistent with all metabolomics analyses in this manuscript. Represented by stacked bar charts are abundance of metabolites enriched (combined ESI modes in Roots (local) and Shoots (systemic) **(e)**.

**Figure S5.**
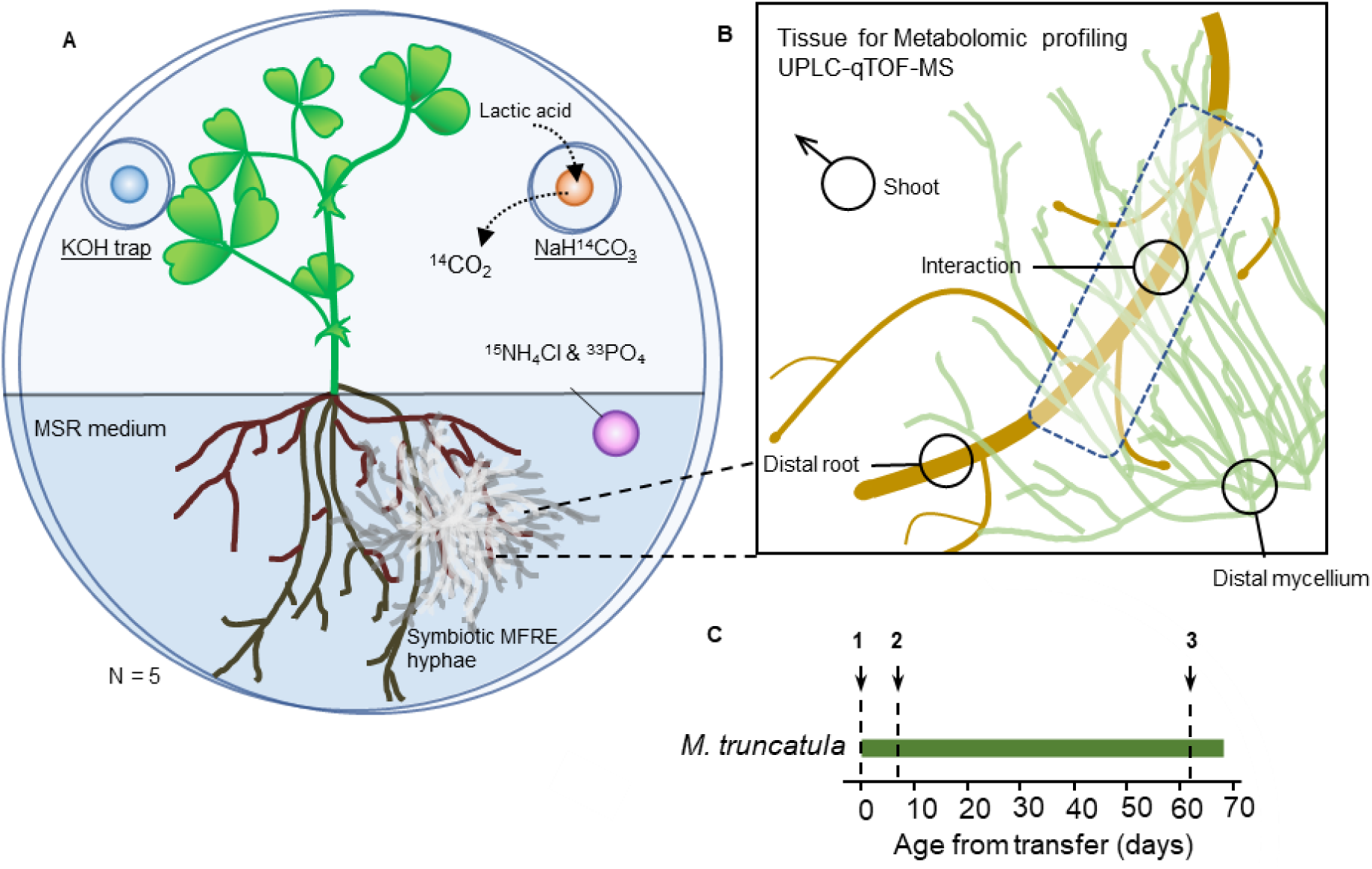
MFRE-Medicago experimental microcosms to quantify nutrient flux between symbionts **(a)**. Microcosms using the same design but without isotope tracers used for metabolomics analyses with location of samples analysed indicated in **(b)**, **(c)** indicates the timing of these experiments where **1** is Scarification and stratification of *M. truncatula* seeds, **2** Seedlings placed on monoxenic plates pre-inoculated with MFRE (above), and **3** Label and harvest root material for microscopy or metabolomics.

---

**Table S1** Adducts from Metlin and mummichog

**Table S2** Volcano output for enriched metabolites and pathways (ESI+ & ESI-). Sheet 1 – Metabolites significantly enriched in symbiotic Roots (local) and Shoots (systemic). Sheet 2 combined annotations from Sheet 1. Metabolites significantly enriched in non-fungal Roots (local) and Shoots (systemic).

**Table S3** mummichog enrichment information

