## Supplementary material for "Common symbiotic signalling pathway not essential for formation of functional mutualisms with endophytic fungi": Methods File 1

### SI Material and Methods

*In vitro culturing of* Medicago *and MFRE isolate*

We used *Medicago truncatula*, cultivar Jemalong in the A17 background, with mutants impaired in AM symbiosis at different genes of the CSSP, namely *dmi2-1* and *dmi3-1*, which dramatically reduce mycorrhization (Oláh et al., 2005), and *ram1-1* which exhibits a near-complete abolition of AM colonisation (Gobbato et al., 2012). *In vitro* axenic cultures of Lyc-1 MFRE-isolate were grown on modified Strullu-Romand (MSR) medium, lacking vitamins and sucrose (Declerck et al., 1998), solidified with 0.4% Gelzan (pH 5.5; PhytoTech, Kansas, US) and used for inoculation. *Medicago* seeds were scarified on sandpaper (120 grit) before surface-sterilisation in bleach for 5 min followed by thorough rinsing in sterilised dH2O before further sterilisation in 70% ethanol for 1 min, and a second rinsing in dH_2_O. Seeds were transferred, under sterile conditions onto MSR medium and left for 3 days in the dark at 4°C. Plates were then transferred under LED lighting and left to germinate.

To establish MFRE-Medicago monoxenic microcosms, MSR medium (above), was plated slanted in 140 mm sterile Petri dishes. 3 mm^2^ plugs of MFRE mycelium were inoculated onto this medium for a week and allowed to establish. Subsequently, 4-day-old *Medicago* seedlings were placed onto the growth medium within 5 mm of establishing MFRE hyphae. Separate control plants were placed in non-inoculated medium. The sterile monoxenic microcosms were sealed with Parafilm, and the root system was covered to limit light entry before being placed vertically and maintained at 22°C, with a 16-h photoperiod and a photosynthetic photon flux of 300 μmol m^−2^s^−1^. After 12 weeks MFRE extraradical mycelium development was observed in *Medicago* cultures and root colonisation was evaluated for intracellular symbiotic structures (Supplementary Fig. 2; Supplementary Fig. 5).

*Colonisation of host plants roots by MFRE*

Approximately 20 mg (fresh weight) of each root system was collected to evaluate MFRE fungal root colonization colonisation. These roots were cleared in 10% KOH for 1 h and stained with 5% ink-vinegar solution for 1 h (Vierheilig et al., 1998). Afterwards, roots were destained in 10% acetic acid for 24 h before being mounted onto slides. The percentage of MFRE root colonisation in terms of presence of intracellular symbiotic structures such as hyphae and spherical, vesicle-like swellings was assessed using the modified gridline intersect method (McGonigle, T. P. et al., 1990) using a compound stereo microscope (GXM-L1500A-HGB; GTVision, Suffolk, UK) at 400 X magnification.

*Quantification of carbon for nutrient exchange between plant and MFRE symbionts*

*In vitro* *Medicago* microcosms were split into two groups: (i) monoxenic cultures of the three *Medicago* mutants and A17 wild-type plants colonised by MFRE (Supplementary Fig. 5; n = 8) and (ii) control plates with A17 Medicago without fungi (n = 5). 30 µL aqueous solution containing 0.25 MBq ^33^P-orthophosphate (111 TBq mmol^−1^ SA, 0.15 ng ^33^P; Hartmann Analytics) and ^15^N-ammonium chloride (2 mg ml^−1^; 0.06 mg ^15^N; Sigma-Aldrich) was introduced into a well dug into the medium of each plate (Supplementary Fig. 5 and see Hoysted *et al.,* 2023). The well was placed at the peripheral edge of the extraradical mycelium, and at a similar distance from the plant roots in no-fungus control plants. Wells were backfilled with Gelzan-MSR medium. This combination of treatments (intact fungal hyphae and non-fungal) allowed us to account for the role of intact MFRE hyphae in the movement of isotopes towards plants within our microcosms. The medium containing *Medicago* roots and MFRE hyphae was sealed from the aboveground plant tissue using a polythene sheet and anhydrous lanolin before each plate was sealed with parafilm and placed in a gas-tight container. A 0.25-MBq pulse of ^14^CO_2_ was liberated into the headspace of sealed Petri dishes by injecting 2 ml of 10% (w/v) lactic acid into containers holding 6.5 μl NaH^14^CO_3_ (specific activity 2.146 GBq mmol^−1^; Revvity; Supplementary Fig. 5) within the chambers. Sealed chambers were maintained for 24 hours under the growth conditions described above.

After 24 hours, plants were carefully removed from the MSR medium, separating roots and shoots, and the medium containing the MFRE extraradical mycelium was collected. All materials were freeze-dried before being weighed and homogenised. For ^33^P, between 0.1 and 2 mg of above-ground tissue (shoot and leaf) was digested in 1 mL concentrated sulphuric acid at 365°C for 15 min. Hydrogen peroxide (50 µL) was added to cooled samples and returned to the digest block for 1 min (BT5D; Grant Instruments, Cambridge, UK). Cleared digests were diluted to 10 mL with dH_2_O. 1 mL of each digest solution was added to 10 mL of Emulsify-safe scintillation cocktail and ^33^P-radioactivity of plant material was quantified through liquid scintillation (Tri-Carb® 4900TR, PerkinElmer, Beaconsfield, UK). Total ^33^P was calculated using the methods described previously (Hoysted et al., 2023) (see SI for details).

Between 0.1 and 2 mg of freeze-dried above-ground tissue were weighed into 6 × 4 mm^2^ tin capsules (Sercon,Crewe, UK), and ^15^N abundance was determined using a continuous flow isotope-ratio mass spectrometer (IRMS; PDZ 2020 IRMS; Sercon) using air as the reference standard; the IRMS detector was regularly calibrated to commercially available reference gases. ^15^N transferred from fungus to plant was calculated using the equations adapted from previous work (Hoysted et al., 2023) (see SI for details). ^14^C activity of shoot, root and fungal samples was quantified through sample oxidation (307 Packard Sample Oxidiser, PerkinElmer, Beaconsfield, UK) followed by liquid scintillation. Total C (^12^C+^14^C) fixed by the plant and transferred to fungus was calculated as a function of the total volume and CO_2_ content of the labelling chamber and the proportion of the supplied ^14^CO_2_ label fixed by plants. The total C budget for each microcosm was calculated using equations adapted from Hoysted *et al.,* 2023 (see SI for details). Total % allocation of plant-fixed C to extraradical symbiotic fungal hyphae was extrapolated from these equations by including the proportion of ^12^C in the system and adding it that of ^14^C. One C sample of the mutant *ram1-1* was lost and is therefore not included in the analysis.

*Total P quantification*

We used colorimetric analysis to determine the total P content of plant roots and shoots. Following acid digestion (see ^33^P determination for method), we measured colorimetric development against a P standard with an adapted version of the molybdate blue reaction (Leake, 1988). Briefly, 0.2 mL of the digest sample was added to a cuvette containing 0.5 mL ammonium molybdate (8 mM) and 0.4 mL ascorbic acid (100 mM) solutions and diluted to 3.8 mL using dH_2_O. P standards of the following concentrations were generated using a 10 mg mL^-1^ of Phosphoric acid; 0.02, 0.05, 0.1,0.2, 0.5 and 1 (µg mL^-1^). Solutions were left in the dark for 45 min before taking measurements at OD_882_ using a spectrophotometer (Jenway 6300). Mass of P in our samples was calculated against the standard curve generated (*y*=8.713, R^2^=0.996) from which the mass of total P in the samples was calculated.

*Metabolomic analysis of plant and fungal tissues*

To determine whether patterns of exchange observed in fungal plants relate to alterations in root metabolic pathways commonly associated with symbiosis (such as carbohydrate biosynthesis), we conducted a micro-scale MS analysis on non-fungal and colonised plant shoot and root material and MFRE extraradical mycelium. Using non-isotope labelled microcosms (Supplementary Fig. 5), we sampled shoots and roots from plants grown in the presence or absence of MFRE extraradical mycelium within the MSR medium from microcosms containing colonised plants. As per the labelling experiment, after 12 weeks plant and fungal tissues were harvested and immediately flash-frozen, before being lyophilised. Samples were homogenised using FastPrep-24 (MP Biomedicals, Derby) prior to chemical solvent extraction in 1 mL of methanol:chloroform:water mix (2.5:1:1 v/v). Samples were vortexed then sonicated for 5 mins (ultrasonic cleaner, VWR) to achieve full cell disruption, before centrifugation at 15,000g for 30 mins (MSE Sanyo Hawk 15/05). 50 µl aliquots were taken into glass mass spectrometry vials (Supelco, Mexico) for untargeted metabolic profiling via UPLC-Q-TOF mass spectrometry (MS). We used the ACQUITY ultra-high-pressure liquid chromatography (UPLC) coupled with SYNAPT G2 Q-TOF mass spectrometer with an electrospray ionization (ESI) source (Waters, UK) at the biOMICS platform (University of Sheffield, UK).

Chromatographic separation was achieved via BEH C18 column (2.1 × 50 mm, 1.7 μm, Waters) with C18 VanGuard pre-column (2.1 x 5 mm, 1.7 µm, Waters) at a flow rate of 0.3 mL min^−1^. The mobile phase consisted of solvent A (0.05 %, formic acid v/v, in water) and solvent B (0.05 % formic acid v/v in acetonitrile) with the following gradient: 0 – 3 min 5 – 35 % B, 3 – 6 min 35 – 100 % B, holding at 100 % B for 2 min, 8 – 10 min, 100 – 5 % B. The column was maintained at 45 °C at an injection volume of 10 μL. After every tenth sample, a QC sample was run (consisting of a pool of every sample in the experiment) and blanks (Solvent A without sample) were injected at the beginning and end of the run to maintain column integrity. Samples were generated in negative and positive ionization mode (ESI- and ESI+) and separated by two consecutive blank injections to stabilise the column between ionization modes. MS detection of ions was operated in sensitivity mode by SYNAPT G2 (50 - 1200 Da, scan time = 0.2 s) for both ionisations, using a full MS scan (*i.e.* no collision energy). The following conditions were applied for ESI-; capillary voltage, 3.15 kV; sampling cone voltage, 10 V; extraction cone voltage, 4.5 V; source temperature, 120 °C; desolvation temperature, 350 °C; desolvation gas flow, 850 L h^-1^; cone gas flow, 105 L h^-1^. Settings were identical for ESI+ apart from a capillary voltage of 3.5 kV. Prior to analyses, the Q-TOF was calibrated by infusing a sodium formate solution. Accurate mass detection was ensured by infusing the internal lock-mass reference peptide leucine enkephalin during each run.

*Data analyses*

Isotope tracing data were analysed in R (ver. 4.4.0). Data were tested for normality and homogeneity of variances using the Kolmogorov–Smirnov test. Where assumptions for parametric tests were not met, data were transformed using log10 (this applied to total C concentration only). The differences between plant assimilation of ^33^P and ^15^N were tested using ANOVA, as were biomass data, colonisation data and ^14^C transfer. Whiskers on box plots represent each of the data points (minimum to maximum) recorded during data collection. For metabolomics data, preprocessing was performed via XCMS (Tautenhahn et al., 2012) using the pipeline described in Parker et al. 2023 (Parker et al., 2023) (https://untargeted-metabolomics-workflow.netlify.app/). Briefly, .RAW files were converted to .mzXML using MSConvert (ProteoWizard (Kessner et al., 2008) before being uploaded to XCMS online (Tautenhahn et al., 2012) to produce a feature table with the HPLC/ Waters TOF parameter ID (#3237). Subsequent data analysis was performed using the MetaboAnalyst platform v3.0(Pang et al., 2020). The tidied data were log-transformed and autoscaled to achieve normality before PCA plots were generated to visualise ordination of the data. Binary comparisons between fungal and non-fungal tissue were performed using Volcano plots with a fold-change of >2 and a p< 0.5. Of the significant metabolite features, metabolite identification was performed using METLIN (Guijas et al., 2018) with tolerance values set below 30 PPM and adducts for ESI- and ESI+ accounted for (see Supplementary Table 1a). Finally, functional pathway analysis utilised the Mummichog2 algorithm against peak intensity data and annotated via the KEGG pathway library for the *Arachis hypogaea* genome (for root analysis) and *Aspergillus niger* genome (for fungal analysis). Shoot comparisons yielded no enhanced pathways with this analysis, so are not presented. Currency metabolites and potential adducts were accounted for (see Supplementary Table 1b) and the P-value cut-off was set at 0.05 and with pathways containing at least 2 entries.

#### Equations, as described in Hoysted *et al.,* 2023

^33^P content was calculated using Eqn [**1**](https://nph.onlinelibrary.wiley.com/doi/full/10.1111/nph.18630#nph18630-disp-0001):

$$M^{33}P = \left\{ \left[ \frac{\frac{\mathrm{cDPM}}{60}}{\mathrm{SAct}} \right]M_{wt} \right\} \mathrm{Df}$$

where *M*^33^P = mass of ^33^P (mg); cDPM = counts as disintegrations per min; SAct = specific activity of the ^33^P course (Bq mmol^−1^); Df = dilution factor; and *M*_wt_ = molecular mass of P

The total C per cent allocation of plant-fixed C to extraradical symbiotic fungal hyphae was calculated by subtracting the activity (in Becquerels) of Medicago cultures without fungus from that detected in monoxenic cultures with MFRE, dividing this by the sum of activity detected in all components of each microcosm (media, roots and shoots), then multiplying by 100. Outlined in Eqn [**2**](https://nph.onlinelibrary.wiley.com/doi/full/10.1111/nph.18630#nph18630-disp-0002):

$$M_{c}=\left( \left( \frac{A}{\mathrm{SAct}} \right)M^{14}C \right)+(P_{r}\times M{wt}_{c})$$

where *M*_c_ = mass of carbon transferred from plant to fungus; *A* = radioactivity of the tissue sample (Bq); SAct = specific activity of the source (Bq Mol^−1^); *M*^14^C = atomic mass of ^14^C; *P_r_* = proportion of the total ^14^C label supplied present in the tissue; 𝑀wt_c_ = mass of C in the CO_2_ present in the labelling chamber (g) (from the ideal gas law; Eqn [**3**](https://nph.onlinelibrary.wiley.com/doi/full/10.1111/nph.18630#nph18630-disp-0003)):

$$M_{\mathrm{cd}}=M_{\mathrm{cd}} \left( \frac{PV_{\mathrm{cd}}}{RT} \right)\therefore m_{c}=m_{\mathrm{cd}}\times0.27292$$

where *m*_cd_ = mass of CO_2_ (g); *M*_cd_= molecular mass of CO_2_ (44.01 g mol^−1^); P = total pressure (kPa); *V*_cd_ = volume of CO_2_ in the chamber (0.003 m^3^); *R* = universal gas constant (J K^−1^ mol^−1^); *T* = absolute temperature (K); *m*_c_ = mass of C in the CO_2_ present in the labelling chamber (g), where 0.27292 is the proportion of C in CO_2_ on a mass fraction basis.
